# ERK1/2 signalling dynamics promote neural differentiation by regulating the polycomb repressive complex

**DOI:** 10.1101/586719

**Authors:** Claudia I. Semprich, Vicki Metzis, Harshil Patel, James Briscoe, Kate G. Storey

## Abstract

Fibroblast Growth Factor (FGF) is a neural inducer in many vertebrate embryos, but how it regulates chromatin organization to coordinate the activation of neural genes is unclear. Moreover, for differentiation to progress FGF signalling has to decline. Why this signalling dynamic is required has not been determined. Here we show that dephosphorylation of the FGF effector kinase ERK1/2 rapidly increases chromatin accessibility at neural genes in mouse embryos and, using ATAC-seq in human embryonic stem cell derived spinal cord precursors, we demonstrate that this occurs across hundreds of neural genes. Importantly, while Erk1/2 inhibition induces precocious neural gene transcription, this step involves dissociation of the polycomb repressive complex from gene loci and takes places independently of subsequent loss of the repressive histone mark H3K27me3 and transcriptional onset. We find that loss of ERK1/2 activity but not its occupancy at neural genes is critical for this mechanism. Moreover, transient ERK1/2 inhibition is sufficient for polycomb protein dissociation and this is not reversed on resumption of ERK1/2 signalling. These data indicate that ERK1/2 signalling maintains polycomb repressive complexes at neural genes, that its decline coordinates their increased accessibility and that this is a directional molecular mechanism, which initiates the process of neural commitment. Furthermore, as the polycomb repressive complexes repress but also ready genes for transcription, these findings suggest that ERK1/2 promotion of these complexes is a rite of passage for subsequent differentiation.

## Introduction

The identities of signals that induce particular cell fates are now well-established, but how such signalling regulates chromatin to coordinate the transcription of differentiation genes and so orchestrate engagement of a differentiation programme is not well understood. Fibroblast growth factor (FGF) signalling has been implicated in the acquisition of neural cell fate in many vertebrate embryos [1–7]; reviewed in [8] (although the timing of involvement varies between species and may reflect differences in induction of anterior and posterior regions [6, 9, 10]). Intriguingly, while FGF is required for neural induction in most of these contexts, its decline is also necessary for differentiation progression.

The requirement for transient FGF signalling to promote neural differentiation is particularly evident in the elongating embryonic body axis in which the spinal cord is generated progressively, as here there is a clear spatial separation of the temporal events of differentiation. FGF acts, along with Wnt signalling, in the caudal lateral epiblast (CLE)/node streak border (which later form the tailbud) to maintain a multipotent cell population known as neuromesodermal progenitors (NMP) which progressively gives rise to the spinal cord and paraxial mesoderm (Figure 1A) [11–15] reviewed in Henrique et al., 2015). Blocking FGF signalling in this cell population accelerates the onset of neural differentiation genes and ectopic maintenance of FGF inhibits this step [16–18]. During normal development differentiation onset is promoted by rising retinoid signalling, provided by adjacent paraxial mesoderm, which represses *Fgf8*, restricting it to the tail end (Figure 1A) [17–20] reviewed in [21]. These findings indicate that decline in FGF signalling promotes neural differentiation, however, little is known about the mechanism(s) by which such signalling dynamics mediate the coordinated activation of neural differentiation genes.

**Figure 1.**
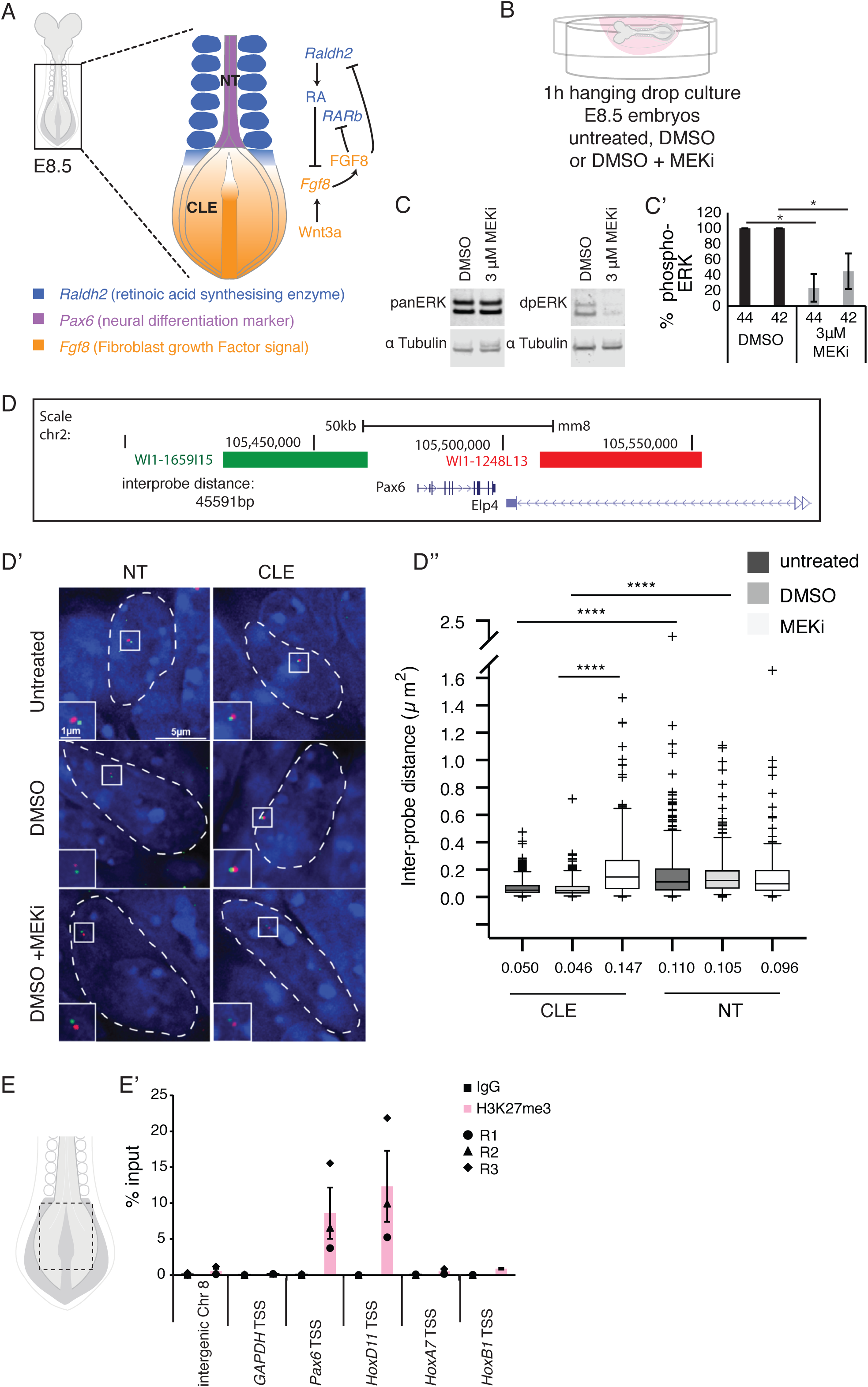
Rapid chromatin decompaction at the neural progenitor gene *Pax6* in the caudal lateral epiblast following ERK1/2 dephosphorylation in the developing mouse embryo. (A) E8.5 mouse embryo schematic, showing the caudal region and signalling pathways known to regulate neural differentiation, NT neural tube, CLE caudal lateral epiblast, RA retinoic acid, *RAR* retinoic acid receptor, FGF, fibroblast growth factor, *Raldh2* Retinaldehyde dehydrogenase 2; (B) Schematic of the hanging drop culture used to treat E8.5 mouse embryos with small molecule MEK inhibitor (MEKi, PD184352) or vehicle control DMSO for 1h; (C and C’) Representative Western blot of embryo lysates probed with antibodies against total (panERK1/2) and phosphorylated ERK1/2 (dpERK1/2) and LiCOR quantification data (n = 3 independent experiments, error bar = SEM, * p= <0.05); (D) Position of fosmid probes flanking the *Pax6* locus (interprobe probe distance ca. 45kb); (D’) representative examples of fosmid probes in individual NT and CLE nuclei (white dashed line) visualised with DAPI (blue) in each untreated condition, vehicle control (DMSO) and MEKi (in DMSO) treated; (D’’) Inter-probe distance measured in > 50 nuclei in NT and CLE in each of three embryos in each condition (n= > 150 nuclei/condition/region, Mann Whitney - test/RankSum-test **** p ≤ 0.0001); (E) Schematic showing caudal end explant (full tissue thickness taken) in E8.5 embryo; (E’) Chromatin from three biological replicates, each consisting of 30 pooled explants, were interrogated for H3K27me3 levels at transcription start sites (TSS) of *Pax6*, known PRC target *HoxD11* and at control regions (see text) compared to IgG background. 3 biological replicates (circle, triangle and diamond) and average (bar) shown, error bar = SEM.

Importantly, as NMP progeny leave the CLE and embark on neural differentiation they experience loss of the FGF effector kinase ERK1/2/MAPK (hereafter referred to as ERK1/2) activity in both chick and mouse embryos [22, 23]. This suggests that ERK1/2 signalling dynamics may regulate onset of neural differentiation. The *in vitro* manipulation of embryonic stem (ES) cells has provided some insight into the involvement of ERK1/2 signalling in this process. Rather than promoting differentiation however, inhibition of ERK1/2 activity in mouse ES cells supports pluripotency [24, 25] and exposure to FGF/ERK1/2 signalling appears here to be an initial step towards differentiation. In agreement with this, FGF is required for neural differentiation of mES cells [26–28] although the timing of this requirement differs between assays [29]. Importantly, however, only a short critical period of ERK1/2 signalling is required in mES cells for subsequent expression of neural genes [27], after which FGF inhibition then accelerates neural differentiation [27, 30, 31]. Moreover, in mouse epiblast stem cells (mEpiSC), which rely on FGF for self-renewal, prolonged FGF signaling abrogates neural differentiation [32]. Consistent with this in both mEpiSCs and human ES cells (which also depend on FGF for self-renewal), inhibition of FGF/ERK1/2 signalling promotes neural differentiation [30, 32]. These findings indicate that temporal control over FGF/ERK1/2 signalling is instrumental in the estabishment of neural identity from epiblast cell precursors.

A clue to the mechanism by which FGF regulates neural differentiation was revealed by analysis of chromatin organisation at key neural progenitor genes in the mouse embryo CLE [33]. This showed that FGF signalling promotes compaction at *Pax6* and further that inhibition of this pathway in *Raldh2* mutant embryos decompacted this locus, even in this retinoid deficient condition in which *Pax6* fails to be expressed [33]. This indicated that FGF regulation of chromatin organisation is molecularly distinct from the machinery that drives subsequent gene transcription, but left open the question of how FGF signalling modifies chromatin organisation. Potential targets for FGF include the polycomb repressive complexes 1 and 2 (PRC1 and 2), which are involved in the repression of differentiation genes in many contexts [34–40] reviewed in [41]. PRC2 is composed of core proteins Ezh2, Eed and Suz12, and via Ezh2 catalyses the methylation of histone 3 lysine 27 (H3K27me2/3) [42–45] and this is augmented in mES cells by the accessory protein, Jarid2 [46–50]. Methylation of H3K27 provides a docking site for PRC1, which in turn directs chromatin compaction and inhibits transcription from target gene loci [51–53] reviewed in [54]. Intriguingly, ERK1/2 inhibition in mES cells, while driving cells into a naïve pluripotent cell state leads to reduction in PRC2 at developmental genes [55–57]. This suggests a role for ERK1/2 signalling in the recruitment or maintenance of polycomb mediated repression. There is also evidence that this may involve regulation by ERK1/2 protein association with PRC occupied chromatin in mES cells [56], although this appears not to be conserved in human ES cells [58].

Here we provide evidence in the caudal mouse embryo and in analogous human ES cell-derived spinal cord precursors, that inhibition of ERK1/2 activity rapidly increases chromatin accessibility at neural differentiation genes. We demonstrate that ERK1/2 dephosphorylation promotes precocious neural gene expression and that loss of PRCs is a molecularly distinct event that precedes transcription. We further show that it is loss of ERK activity, but not its association with chromatin, that leads to the disassociation of polycomb proteins from neural gene loci. Strikingly, this loss of PRCs requires only transient ERK1/2 inhibition and is not reversible upon resumption of ERK1/2 signalling. These findings reveal the succession of genome-wide, molecular mechanisms operating downstream of FGF/ERK1/2 signalling, that engage the neural differentiation programme.

## Results

### Chromatin compaction around neural differentiation gene *Pax6* in mouse embryos is ERK1/2 dependent

To determine whether ERK1/2 signalling regulates chromatin organisation at neural differentiation gene loci *in vivo*, we exposed whole mouse embryos (E8.5) to the small molecule MEK inhibitor PD184352 (MEKi) or vehicle only DMSO control for one hour (Figure 1B). The efficiency of ERK1/2 inhibition was determined by quantifying levels of phosphorylated ERK1/21/2 (42 and 44 kD) in whole protein extracts from MEKi and DMSO treated embryos (Figures 1C, C’). This regime was then used to measure chromatin compaction around the *Pax6* locus using Fluorescent *In Situ* Hybridisation (FISH) with fosmid probes hybridising either side of the locus [33] (Figure 1D). Chromatin compaction was assessed by measuring inter-probe distance in *Pax6* transcribing neural tube cells and in CLE cells where *Pax6* has yet to be expressed (Figures 1D’, D’’). In untreated embryos, the inter-probe distance was greater in neural tube than in CLE nuclei (Figures 1D’, D’’). Similar observations were made in the DMSO treated embryos (Figures 1D’, D’’) and these two control conditions were not significantly different, suggesting that DMSO did not alter chromatin organisation. By contrast, the *Pax6* locus was more open in CLE nuclei of MEKi treated embryos compared to both controls (Figures 1D’, D’’). This decompaction correlated with a 1.7 fold decrease in the number of base pairs per nm compared to controls. These findings indicate that ERK1/2 activity promotes chromatin compaction at this neural differentiation gene locus and that active ERK1/2 is acutely required for this action *in vivo*.

### Polycomb-mediated histone modification H3K27me3 marks the neural differentiation gene *Pax6 in vivo*

To determine whether the PRCs regulate chromatin compaction around the *Pax6* locus *in vivo*, we carried out chromatin immunoprecipitations (ChIPs) for the PRC-mediated histone modification H3K27me3. Mouse embryo microdissection was performed to enrich for the caudal region at E8.5 for ChIP-qPCR (Figure 1E and see Methods). The *Pax6* transcription start site (TSS) was marked by high levels of H3K27me3 indicative of polycomb activity and consistent with repression of this gene in the CLE (Figure 1E’). Similar levels were also detected at the known PRC target *HoxD11* [59–61], which is also not expressed in the mouse CLE at E8.5 [62]. By contrast, known PRC targets transcribed in the CLE, *HoxA7* and *HoxB1*, [63], exhibited minimal levels of H3K27me3 at their TSSs. This was similar to levels detected at control intergenic region and house-keeping gene *GAPDH* indicative of a lack of polycomb activity at these loci (Figure 1 E’). Together these data demonstrate that polycomb activity at target genes is selective *in vivo*, and identify *Pax6* as a potential polycomb target. This raises the question of whether ERK1/2 activity directs polycomb occupancy and chromatin compaction and thus determines the onset of neural gene transcription.

### *PAX6* locus decompaction correlates with polycomb protein dissociation, H3K27me3 loss and transcription onset in a human *in vitro* model of spinal cord differentiation

To test this, we developed an *in vitro* system to study neural differentiation during the generation of the spinal cord. We reasoned that the use of human ESCs, with the slower progression to neural progenitor cell identity, would provide a better temporal resolution of mechanisms mediating neural differentiation. To this end, we directed the differentiation of human ESCs towards spinal cord [13, 64] (Figure 2A). This regime involved the generation of NMP-like cells on day 3 (NMP-L D3), characterised by the co-expression of *BRA* and *SOX2* and their differentiation into *PAX6* expressing neural progenitors by day 8 (NP D8), which also transcribed the known PRC target *HOXD11* (Figures B-B’’’). More detailed analysis further revealed low level onset of *PAX6* on D6, anticipating the robust expression detected on D8 (Figure 2B’’’). We next asked whether changes in chromatin compaction at *Pax6* occur during human neural differentiation *in vitro*, as we had observed in mouse *in vivo* (Figure 1D). FISH was carried out in NMP-L (D3) and NPs (D8) cells and the distances between the *PAX6*-flanking probes were measured (Figures 2C-C’’). This revealed a significant increase in inter-probe distance in NPs compared to NMP-Ls (Figures 2C, C’). This indicates that the regulation of chromatin compaction is a conserved mechanism operating during neural differentiation, that can be examined and recapitulated *in vitro*.

**Figure 2.**
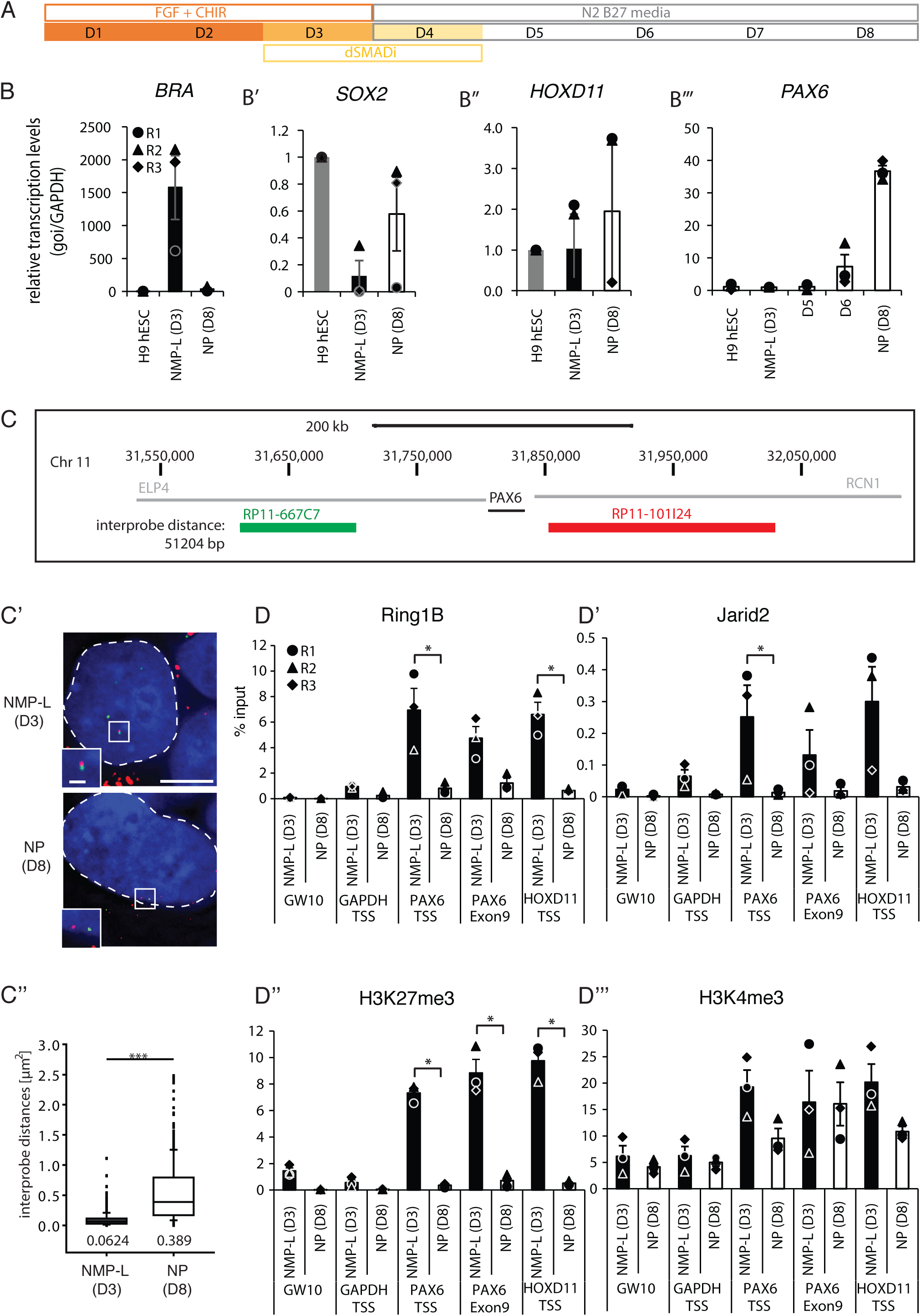
*In vitro* differentiation of human ESC-derived NMP-L cells to NPs reveals locus decompaction correlates with dissociation of polycomb repressive complexes and transcriptional onset at *PAX6*. (A) Schematic of differentiation regime used to generate first NMP-like cells (NMP-L (D3)) and then spinal cord progenitors (NPs (D8)) from human ESCs (D1 = Day1); (B-B’’’) transcription levels of *BRA, SOX2, HOXD11* and *PAX6* assessed by RTqPCR in undifferentiated cells (hESCs), NMP-Ls (D3) and NPs (D8) for *PAX6* additionally on day 5 (D5) and day 6 (D6) of differentiation (n=3 independent experiments, error bars = SEM). (C-C’’) FISH to assess chromatin compaction around the *PAX6* locus in NMP-L and NP. 2 probes flanking the target locus (inter-probe distance ca. 51 kb) were hybridised and labelling visualised in DAPI stained nuclei (blue, outlined with white dashed line). Inter-probe distance measurements in >50 nuclei in NMP-Ls and NPs in 3 individual experiments (n= > 150 nuclei/cell type, Mann Whitney - test/RankSum-test, *** p ≤ 0.001). (D-D’’’) ChIP-qPCRs investigating polycomb repressive complex occupancy (Ring1B/PRC1 and Jarid2/PRC2) and the histone modifications H3K27me3 and H3K4me3 in NMP-L and NP cells (n = 3 independent experiments indicated with circles, triangles and diamonds, bar = average, error bar = SEM, * p≤ 0.01, t test).

To elucidate how chromatin compaction is regulated at the *PAX6* locus, we interrogated PRC occupancy over developmental time. PRC1/(Ring1B) and PRC2/(Jarid2) occupancy, along with H3K27me3 were analysed by ChIP-qPCR at *PAX6* and control loci in NMP-Ls and NPs. These analyses established the presence of PRC proteins and H3K27me3 at the TSS and along the gene body of *PAX6* and at the TSS of known PRC target *HOXD11* in NMP-L cells and importantly demonstrated reduced occupancy at *PAX6* and *HOXD11* in NPs (while baseline detection in the gene desert region (*Gw10*) and the housekeeping gene *GAPDH* remained unchanged) (Figures 2D-D’’). Strikingly, analysis of the active histone modification H3K4me3 revealed no change at the *HOXD11* or *PAX6* TSSs during neural differentiation (Figure 2D’’’). This suggests that polycomb mediated repression is the major activity restricting *PAX6* gene expression in the NMP-L cells, not the low levels of H3K4me3. Together these findings demonstrate that this site-specific deposition of PRC is transient and that its removal is characteristic of the neural progenitor cell state.

### Progressive global increase in neural gene accessibility during *in vitro* human differentiation

While these data suggested that PRC loss is required for the onset of neural gene transcription, it remained unclear whether this regulation of chromatin accessibility is specific for *PAX6* or reflects a general mechanism that mediates onset of neural gene transcription. To address this we performed a global analysis of the chromatin accessibility landscape in cells as they differentiated into NPs from the NMPL cell state, using ATAC-seq [65]. This involved differentiation of NMPL (D3) cells (as in Figure 2A) and sampling cell populations at intervals (D5, D6 and D8). Analysis of the chromatin configuration across the *PAX6* locus revealed increased accessibility at multiple sites, some as early as D5 and most prominent by D6 (Figure 3A), including one region described as an intragenic *PAX6* enhancer region [66] (red box Figure 3A). These early increases in chromatin accessibility correlated with decline in PRC protein occupancy at the *PAX6* locus detected by ChIP-qPCR during differentiation from D5 to D7 (Figures 3B, B’), which becomes significant by D8 (Figures 2D, D’). In contrast with this apparently progressive decrease, particularly in Jarid2 occupancy, H3K27me3 levels at the *PAX6* TSS and along the gene body remain unchanged between D5 and D7 (Figure 3B’) and drop dramatically only at D8 (Figure 2D’’). These data align decreasing PRC occupancy with increasing chromatin accessibility, while acute loss of H3K27me3 appears a later step that coincides with robust *PAX6* transcription.

**Figure 3.**
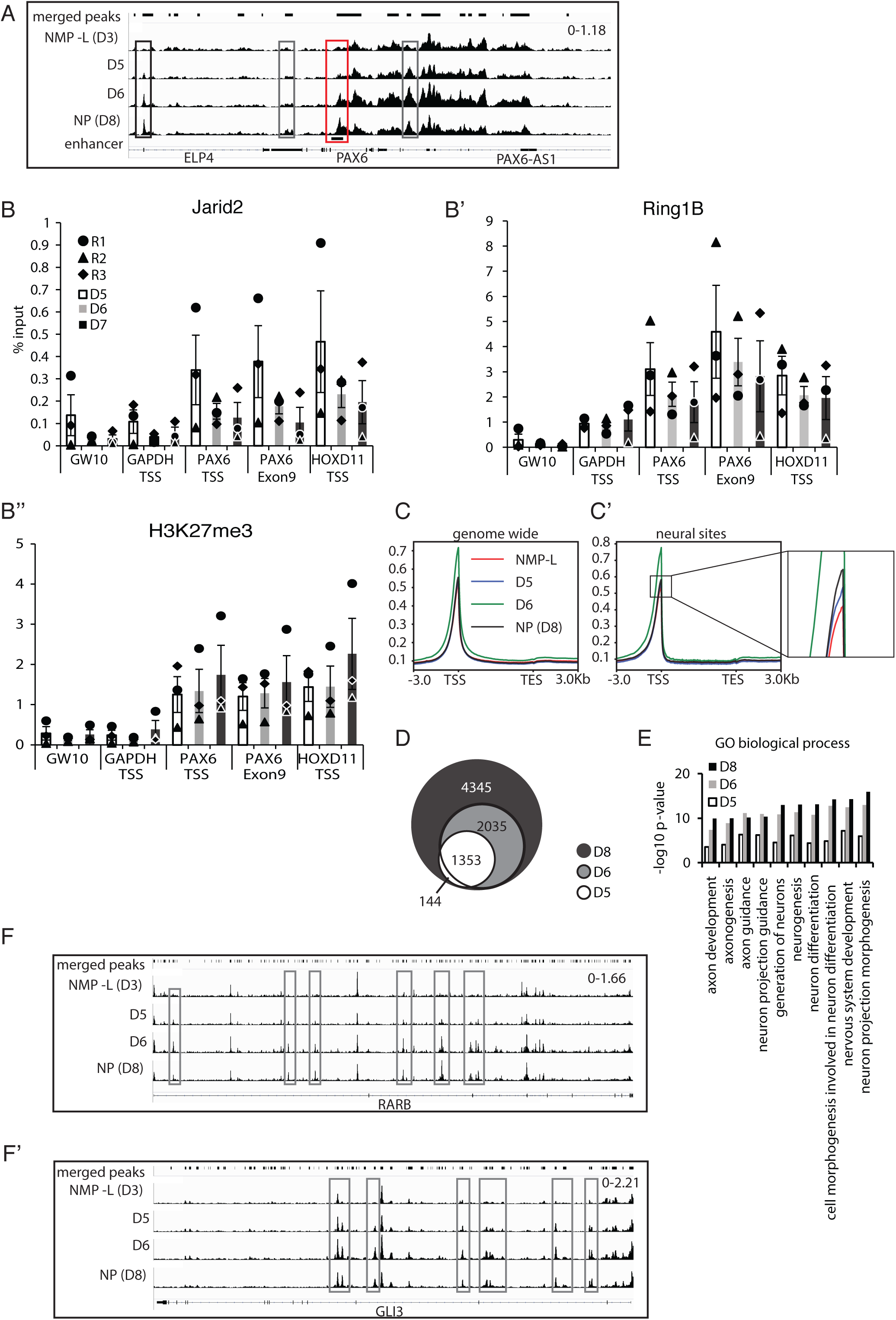
Progressive increase in human neural gene accessibility during *in vitro* differentiation. (A) ATAC-seq peak tracks in NMP-L (D3), D5, D6 and NP (D8) cells for the genomic regions of *PAX6*, grey boxes are peaks within the gene body, black boxes, peaks outside of the gene body and red box genomic region of a known *PAX6* enhancer; (B-B’’) ChIP-qPCR for Jarid2, Ring1B and H3K27me3on D5, D6 and D7 (n = 3 independent experiments, each indicated by circles, triangles and diamonds, bar = average, error bars = SEM, no significant differences between samples, t-test). (C-C’) Metaprofiles comparing chromatin accessibility in NMP-L (D3) and NP (D8) cells along the gene body genome-wide (C) and focussed on genes associated with increased accessibility in NPs (D8) (neural sites, C’) compared to NMP-L; (D) Venn diagram representing distribution of genomic regions more accessible in NPs (D8) (neural sites) compared to NMP-L (D3) and their appearance over time (D5 and D6). Comparison of accessible regions and associated genes on D5 and after D5, revealed ~19% of genes had additional accessible regions, while the majority were found in newly accessible genes, similarly 40% of D6 genes with additional regions became more accessible in NP (D8); (E) GO term analysis of genes associated with neural sites compared to NMP-L (D3) and genes associated with more accessible chromatin regions on D5 and D6 out of the neural sites; (F-F’) ATAC-seq peak tracks in NMP-L (D3), D5, D6 and NP (D8) cells for the genomic regions of *RARB* (E) and *GLI3* (E’), grey boxes are peaks within the gene body

Genome-wide comparison of accessibility patterns across TSSs and gene bodies in D3, D6 and D8 cells revealed that although global accessibility levels remained relatively constant over time, a clear peak in accessibility appeared at D6 of this differentiation (Figure 3C). Detailed analysis comparing D3 and D8 identified 7877 regions with increased, and 11603 regions with decreased, accessibility (analysis performed with diffReps with thresholds FDR ≤ 0.01 and log2FC ≥1). These regions of increased accessibility correlate well with known active enhancer marks such as H3K4me1 and H3K27ac detected in human embryonic spinal cord and brain (data from the ENCODE regulatory element database [67](see Figure S1) and we termed these regions “neural sites”. Analysis of these 7877 neural sites revealed that they are associated with 4001 genes (Table S1) and that accessibility increased across the TSS and body of these genes over time, with accessibility peaking at D6 (Figure 3C’) – this included additional chromatin opening across genes already accessible at D3 as well as new genes with increased accessibility by D8 (Figure 3D).

GO term analysis of the genes associated with regions of increased accessibility on days 5, 6 and 8 revealed terms related to neural development and showed a progression with more genes associated with these terms over time (Figure 3E, Table S1). Focussing on the GO term Neurogenesis we selected 2 key neural progenitor genes in addition to *PAX6* and examined the ATAC-seq signal in detail around the gene body. Chromatin accessibility around *RARB* and *GLI3* showed an increase similar to that associated with *PAX6* as early as D5 (Figures 3F, F’). Overall, these findings indicate that coordinated regulation of chromatin accessibility is a general mechanism that mediates the onset of neural gene transcription, while analysis of PRC occupancy dynamics at the exemplar *PAX6* locus identifies this repression complex as a target of such regulation.

### ERK1/2 dephosphorylation induces precocious polycomb protein dissociation and global increase in chromatin accessibility at neural sites

In chick and mouse embryos, FGFR inhibition elicits precocious *Pax6* transcription [16, 33] and FGFR and ERK1/2 dephosphorylation (this study) decompact this gene locus. We therefore next tested whether ERK1/2 signalling regulates polycomb protein occupancy and if such signalling globally directs chromatin accessibility at our defined neural sites. To address this, we first tested whether inhibition of ERK1/2 signalling advances human neural differentiation *in vitro*; cells were cultured as in Figure 4A and exposed to MEKi or DMSO cells during differentiation (Figure 4B) and *PAX6* transcript levels analysed by RT-qPCR. This revealed precocious *PAX6* transcription beginning at D5 and peaking at D6 in the presence of MEKi (Figure 4C). This correlated with changes in polycomb protein occupancy: MEKi exposure on D5 and D6 resulted in decreased Ring1B at the *PAX6* and control locus *HOXD11*, and Jarid2 levels also showed a declining trend (Figures 4D, D’ and supplementary Figure S2). Furthermore, reduced H3K27me3 levels at the *PAX6* locus on D6 (Figure 4D’’) correlated with maximum precocious *PAX6* transcription elicited by inhibition of ERK1/2 signalling (Figure 4C).

**Figure 4.**
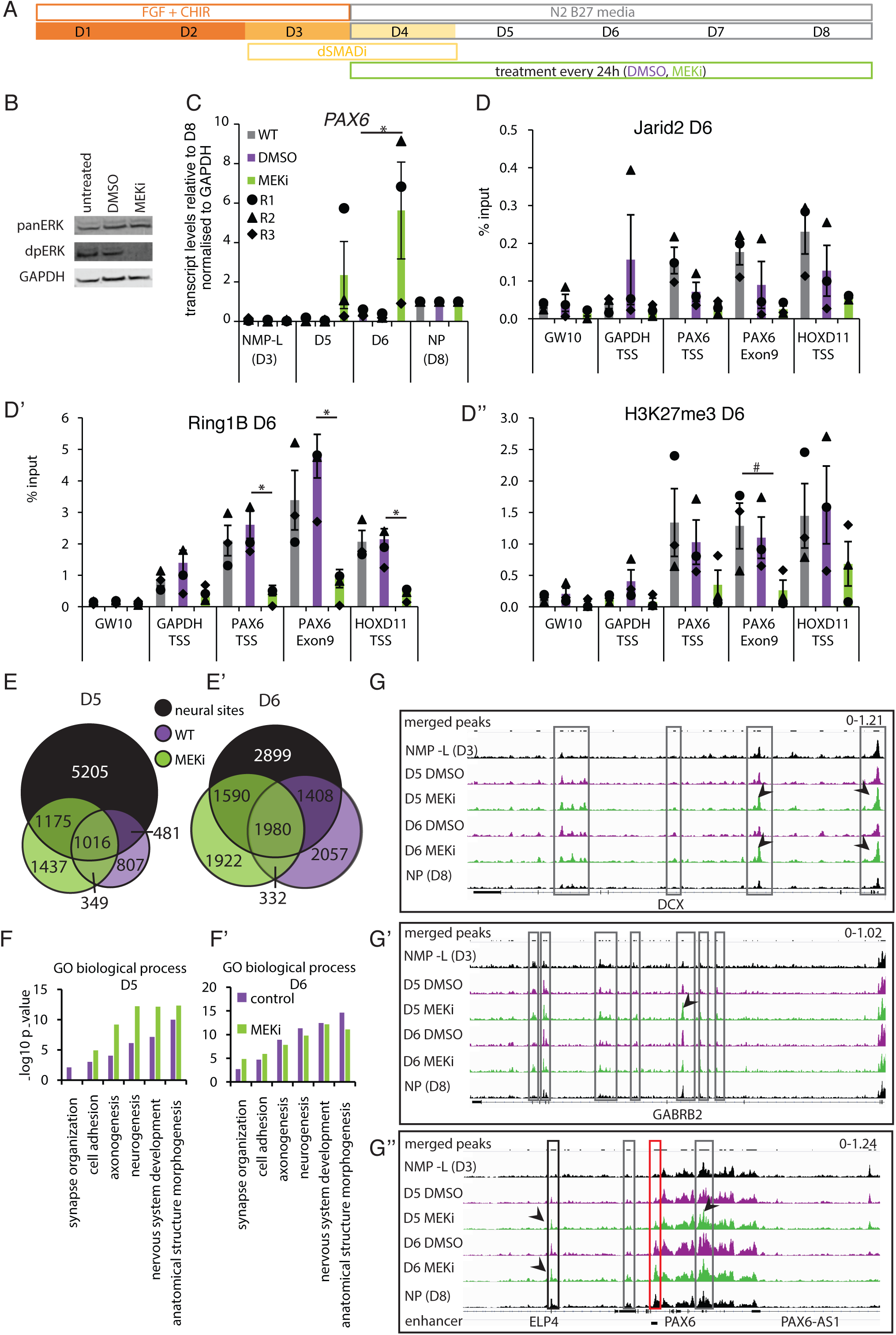
ERK1/2 de-phosphorylation induces precocious *PAX6* expression, reduced polycomb protein occupancy and H3K27me3 at this locus and increases accessibility of hundreds of neural differentiation genes. (A) Differentiation protocol generating NMP-L (D3) cells and spinal cord NPs by D8 and additional treatment with vehicle control DMSO or MEK inhibitor (PD184352, MEKi) every 24h during this differentiation; (B) Western blot confirming that exposure to MEKi lead to reduced ERK1/2 phosphorylation after 24h in differentiation conditions (D3-D4); (C) *PAX6* transcript levels in NMP-L (D3), in control conditions (WT and DMSO) or in MEKi treated cells on D5 and D6 and NP(D8) determined by RTqPCR (n = 3 independent experiments indicated by circles, triangles and diamonds, bar = average, error bars = SEM); (D-D’’) ChIP-qPCR for Jarid2 and Ring1B and H3K27me3 on D6 (n = 3 independent experiments indicated with circles, triangles and diamonds, bar = average, error bars = SEM, * = p ≤ 0.05, t-test); (E-E’) Venn diagrams representing distribution of genomic regions that become more accessible on D5 (E) and D6 (E’) in control condition and MEKi treated cells compared to NMP-L (not restricted to regions accessible in NPs (D8)) as well as more accessible regions on D8; (note comparing DMSO treated cells with untreated cells revealed that these two conditions did not differ from each other on D5 or D6 (Figure S2); (F-F’) GO term analysis of genes associated with more accessible chromatin in D5 (F) and D6 (F’) in control condition and MEKi treated cells compared to NMP-L; (G-G’’) ATAC-seq peak tracks in NMP-L, D5 DMSO and MEKi, D6 DMSO and MEKi and NP (D8) cells for the genomic regions of *DCX* (G), *GABRB2* (G’) and *PAX6* (G’’), grey boxes outline peaks within the gene body, black boxes peaks outside of the gene body and red box genomic region of a known *PAX6* enhancer, example comparisons between peaks in MEKi and DMSO conditions indicated with black arrowhead pairs.

To determine whether ERK1/2 activity acts globally to regulate chromatin accessibility around neural differentiation genes we performed ATAC-seq on cells at D5 and D6 exposed to MEKi or DMSO during differentiation. A comparison of all regions opening between NMP-L (D3) and D5 in control (2653 regions) and MEKi (3977 regions, associated with 417 newly-accessible genes, and 368 more accessible genes) conditions, identified 2612 regions with additional chromatin accessibility following MEKi treatment (Figure 4E, Table S2). Most of the MEKi induced regions of increased accessibility (2191, 55%) belonged to the previously defined neural sites (Figure 3C’), while a further 1437 regions were unique to MEKi treatment and not included in neural sites (Figure 4E). Analysis of D6 control and MEKi datasets revealed a similar pattern, with 3512 additional regions of increased accessibility induced by MEKi (associated with 593 newly accessible genes and 437 genes acquiring additional accessibility) (Figure 4E’) and while the majority coincide with neural sites, a further 1992 regions were detected outside of control D6 and D8 data sets (Figure 4E’ and see below).

To interpret the genome-wide chromatin accessibility changes induced by MEKi, we analysed the GO biological processes associated with these gene cohorts. Strikingly, MEKi treatment on D5 corresponded to an increased representation of genes associated with key neural development terms (Figure 4F). Moreover, at D6 of MEKi treatment, genes associated with later neural development, such as synapse organisation, were specifically increased (Figure 4F’). We therefore analysed the GO terms for genes associated with more accessible regions that were unique to the MEKi data sets on D5 and D6 and found a strong bias towards later neuronal differentiation (Table S3). This suggests that MEKi treatment leads to increased global accessibility at neural differentiation genes and the accelerated progression to a more advanced differentiation cell state. Examining the ATAC-seq peaks at individual genes (*DCX, GABRB2* and *PAX6*) supported the conclusions of the global ATAC-seq analyses, showing that peaks increased over time and that this was advanced with MEKi treatment on D5 (arrowheads in Figures 4G-G”).

### De-phosphorylation does not alter ERK1/2 protein association with chromatin

To investigate the mechanism by which ERK1/2 signalling maintains PRC occupancy at differentiation genes we asked whether ERK1/2 acts directly to mediate recruitment of polycomb machinery, as reported in mES cells [56]. ChIP-qPCR was first used to determine ERK1/2 occupancy across *PAX6* and control genes during in vitro differentiation. In NMP-L cells, ERK1/2 was found at the *PAX6* TSS and gene body and the *HOXD11* TSS and was also detected at the *GAPDH* TSS. Although ERK1/2 localisation at both *PAX6* and *HOXD11* decreased in a manner consistent with increased accessibility of these loci during differentiation, this also occurred at the locus of the constitutively transcribed gene *GAPDH* (Figure 5A). We next analysed whether ERK1/2 occupancy is dependent on its phosphorylation status by treating cells with MEKi during differentiation (as in Figures 4A). No statistical difference was found between MEKi treated cells and control (untreated or DMSO treated) on either D5 or D6 (Figures 5B, B’). These data reveal no consistent relationship between ERK1/2 chromatin association and gene transcription (ERK1/2 is found at the constitutively expressed *GAPDH* locus as well as *PAX6* and other loci) during differentiation. Moreover, the pattern of ERK1/2 occupancy is unaltered upon its de-phosphorylation indicating that change in ERK1/2 activity, but not its chromatin association, regulates PRC occupancy.

**Figure 5.**
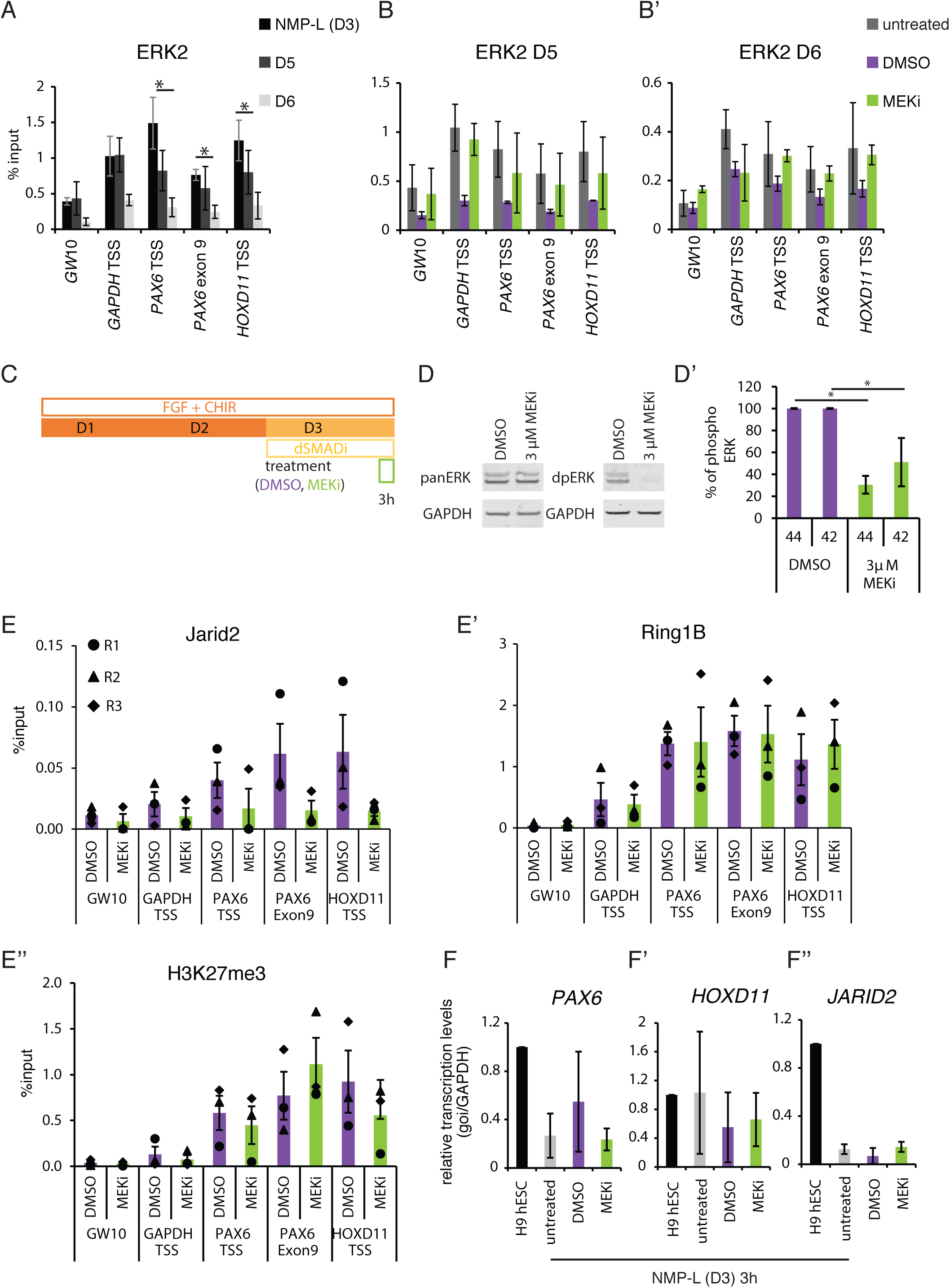
ERK2-chromatin association does not correlate with PRC occupancy and is not regulated by ERK1/2 activity; short ERK1/2 dephosphorylation in NMP-L cells attenuates PRC2 occupancy at the *PAX6* locus, without affecting PRC1 occupancy, H3K27me3 levels or transcription. (A) ChIP-qPCR detecting ERK1/2 occupancy at *PAX6* and control loci during differentiation (NMP-L (D3), D5 and D6, black, dark grey and light grey respectively) (n = 3 independent experiments, error bars = SEM, * = p ≤ 0.05, # = p= 0.055 t-test); (B and B’) ChIP-qPCR investigating ERK1/2 occupancy at the *PAX6* locus on D5 and D6 of the differentiation protocol comparing untreated, DMSO and MEKi exposed samples (n = 3 independent experiment, error bars = SEM, no significant differences between treatments, t-test); (C) Differentiation protocol used to generate NMP-L (D3) cells and treatment regime with vehicle control DMSO or MEKi for 3h; (D-D’) representative Western blot of cell lysates probed with antibodies against total (panERK1/2) and dual-phosphorylated-ERK1/2 (dpERK1/2) and LiCOR quantification data (n = 3 independent experiments, error bar = SEM, * p= <0.05); (E-E’’) ChIP-qPCRs investigating Jarid2 and Ring1B occupancy and H3K27me3 levels at *PAX6* and control regions in NMP-L (D3) cells treated with MEKi or DMSO for 3h (n = 3 individual experiments indicated with circles, triangles and diamonds, bar = average, no significant differences between samples, t-test); (F-F’’) transcription levels of *PAX6, HOXD11*, and *JARID2* assessed by RTqPCR in undifferentiated cell (hESCs), untreated, vehicle control (DMSO) treated or MEKi treated NMP-L (D3) cells (n=3 individual experiments, no significant differences between samples, t-test).

### Short duration ERK1/2 dephosphorylation in NMP-L cells suggests PRC2 loss initiates chromatin changes

To uncover how ERK1/2 activity regulates PRC occupancy we next analysed the timing of Jarid2, Ring1B and H3K27me3 loss from key loci following exposure of NMP-L (D3) cells to MEKi or DMSO (Figure 5C). After only 3 hours, MEKi treatment (in the presence of FGF) reduced dpERK1/2 levels (Figures 5D, D’). Importantly, under these conditions ChIP-qPCR revealed no change in Ring1B nor H3K27me3 occupancy (Figures 5E’ and E’’). However, there was a marked decrease in Jarid2 at *Pax6* and *HoxD11* loci (while little H3K27me3, Ring1B and Jarid2 were detected at negative control loci in any condition) (Figure 5E). This short exposure is therefore insufficient to disassemble the PRC1 (Ring1B) complex or remove the H3K27me3 mark, but may initiate the departure of the PRC2 component Jarid2. Interestingly, no effects on *PAX6* or *HOXD11* transcription, nor reduction in *JARID2* transcription were observed (Figures 3D-D’’). These data raise the possibility that ERK1/2 activity regulates PRCs by supporting Jarid2 occupancy and suggest that Jarid2 dissociation is an initial step in the disassembly of this complex, distinct from later mechanism(s) regulating transcriptional onset of target genes.

### Transient ERK1/2 dephosphorylation in NMP-L cells induces dissociation of both Jarid2 and Ring1b and chromatin decompaction at the *PAX6* locus, but does alter H3K27Me3 nor elicit transcription

To assess the consequences of loss of ERK1/2 activity at a later time point in NMP-L (D3) cells, these were exposed to MEKi for 12 hours before conducting the same panel of ChIP-qPCR experiments (Figure 6A). At the end of this period ERK1/2 phosphorylation levels had now returned to control levels (Figures 6B, B’, indicative of transient MEKi effectiveness), although other consequences of reduced ERK1/2 activity, such as increased PKB phosphorylation were still apparent (Figure S3): this regime therefore delivers only transient inhibition of ERK1/2 activity for at least 3h. However, after 12h reduced Jarid2 and also Ring1B occupancy were now detected at *PAX6* and *HOXD11* loci in exposed cells (while control genomic regions possessed only low level H3K27me3 and Ring1B and Jarid2, which did not change with MEKi treatment)(Figure 6C-C’). Importantly, none of the gene loci analysed exhibited change in H3K27me3 levels (Figure 6C’’) nor did the loss of both PRC1 and PRC2 proteins correlate with transcription of *PAX6* or *HOXD11* nor with reduced *JARID2* transcripts in these conditions (Figures 6D-D’’).

**Figure 6.**
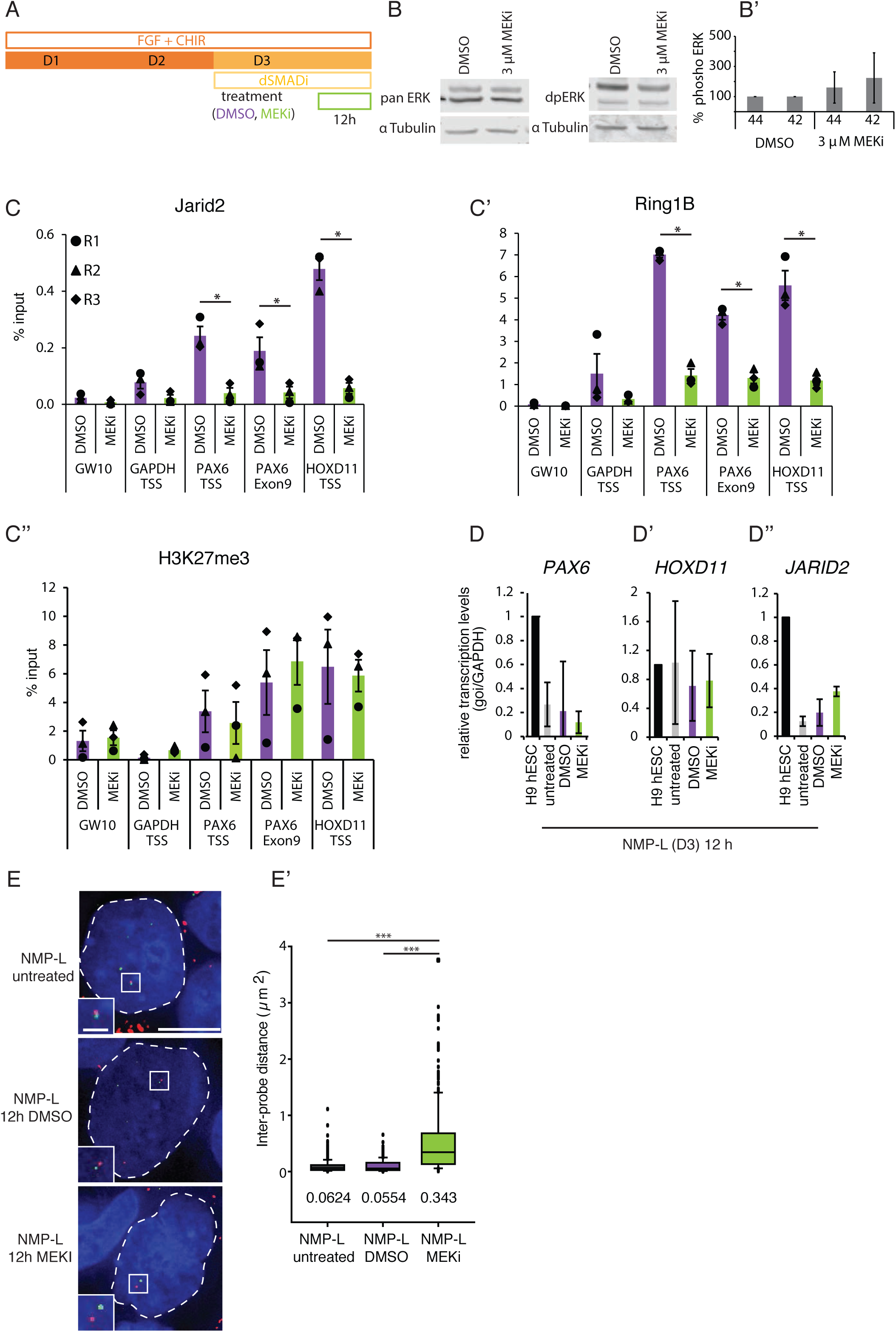
Transient ERK1/2 dephosphorylation induces dissociation of Jarid2 and Ring1B and chromatin decompaction in NMP-L cells at the *PAX6* locus, but does not reduce H3K27me3 nor elicit transcription. (A) Differentiation protocol used to generate NMP-L (D3) cells and additional treatment with vehicle control DMSO or MEKi for 12h; (B-B’) representative Western blot of cell lysates probed with antibodies against total (panERK1/2) and dual-phosphorylated-ERK1/2 (dpERK1/2) and LiCOR quantification data (n = 3 independent experiments, error bar = SEM); (C-C’’) ChIP-qPCRs investigating the histone modification H3K27me3 and polycomb repressive complex occupancy in NMP-L (D3) cells treated with MEKi or DMSO for 12h (n = 3 individual experiments indicated with circles, triangles and diamonds, bar = average, * = p≤ 0.05, t-test); (D-D’’) Transcription levels of *PAX6, HOXD11*, and *JARID2* assessed by RT-qPCR in undifferentiated cell (hESCs), NMP-L cells untreated, vehicle control (DMSO) treated or MEKi treated (n=3 individual experiments, no significant differences between samples, t-test); (E-E’) FISH to assess chromatin compaction around the *PAX6* locus in NMP-Ls untreated and treated with DMSO or MEKi for 12h. 2 probes flanking the target locus (inter-probe distance ca 51 kb) were hybridised and visualised by differential labelling nuclei (outlined with white dashed line) visualised with DAPI (blue). Inter-probe distance measurements in >50 nuclei in the three conditions in three individual experiments (n = > 150 nuclei/cell type, Mann Whitney - test/RankSum-test, *** p ≤ 0.001), this decompaction correlated with a 2.5 fold decrease in number of base pairs per nm compared to both controls (untreated NMP-L (D3): 205 bp/nm, DMSO NMP-L (D3): 218 bp/nm and MEKi NMPL (D3): 87 bp/nm).

These data indicate that a transient reduction in ERK1/2 signalling is sufficient to trigger loss of both PRC2 and PRC1 proteins. To determine the significance of such loss in this context, we further assessed whether 12h MEKi exposure also altered chromatin accessibility around the *PAX6* locus using FISH. Significant decompaction of this region was found in MEKi treated cells compared with both DMSO and untreated controls (Figures 6E, E’). These findings correlate PRC loss with a distinct increase in chromatin accessibility across this key neural progenitor gene and in showing that this step is not reinstated on resumption of ERK1/2 activity, suggest that this is a directional molecular mechanism. Moreover, these data demonstrate that removal of H3K27me3 is not required for such chromatin re-organisation and indicate that regulation of this histone modification is molecularly distinct from the initial effects of ERK1/2 dephosphorylation, indeed this correlated well with later transcriptional onset (Figures 6C −C’’ compare Figures 2 D-D’’). These distinct steps are summarised in Figure 7.

**Figure 7.**
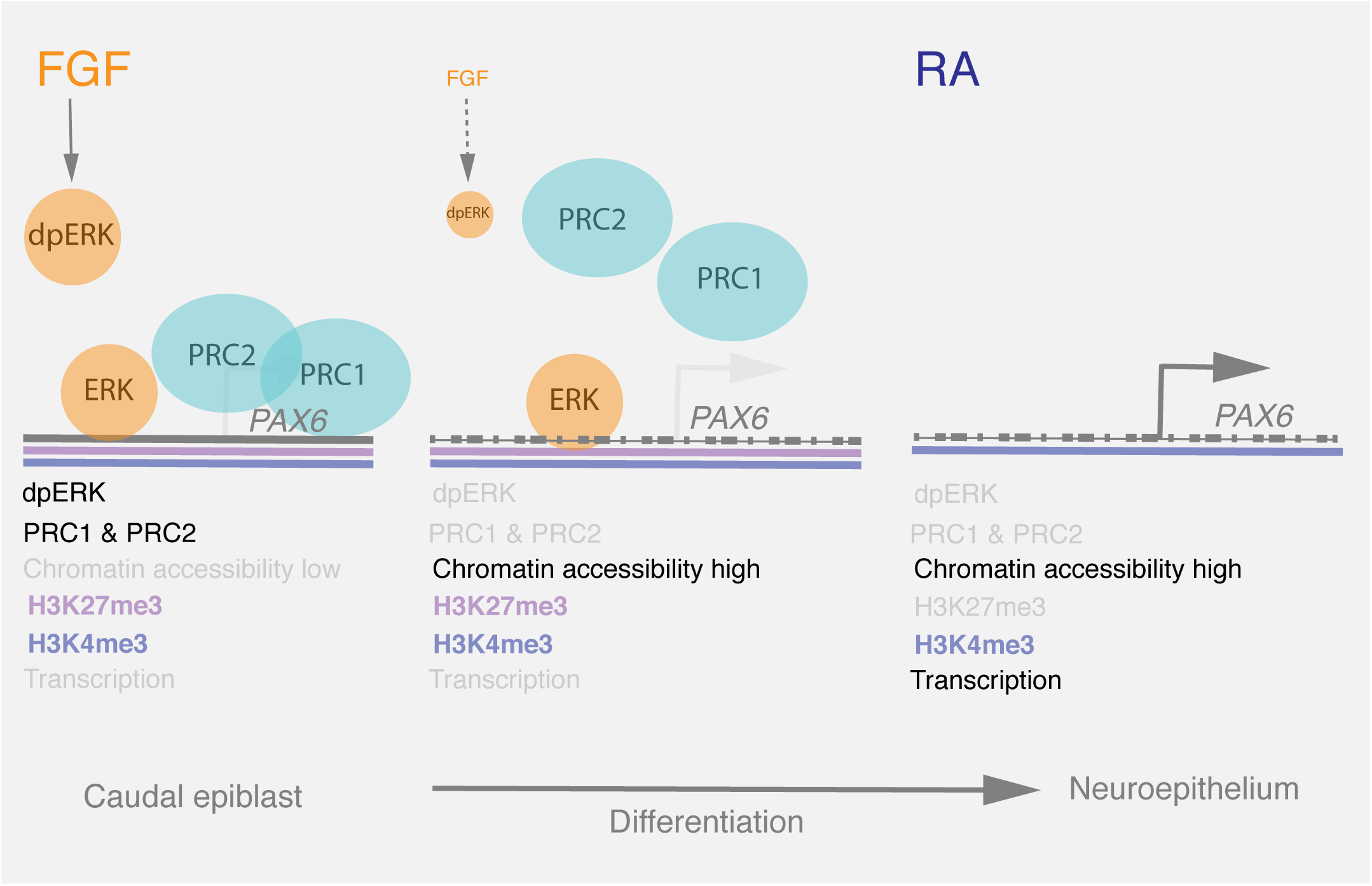
Summary model. Decline in Fibroblast Growth Factor (FGF) and downstream effector kinase ERK1/2 signalling as caudal epiblast cells commence differentiation leads to loss of polycomb repressive complexes (PRC2 and PRC1) from exemplar neural differentiation gene *PAX6* and chromatin decompaction across this gene locus: such reduction in ERK1/2 activity induces genome-wide increase in chromatin accessibility across neural genes. Histone modifications H3K4me3 and H3K27me3 are both detected (bivalent configuration) in the presence of FGF and following dissociation of PRCs, and the PRC-mediated gene silencing mark H3K27me3 is lost only later coincident with transcriptional onset: this indicates that this is a distinct regulatory step, which may depend on retinoic acid (RA) signalling which is known to be required for *PAX6* transcription. As transient loss of ERK1/2 activity is sufficient to remove PRC2 and PRC1 and these complexes are not re-imposed on resumption of ERK1/2 signalling, regulation of ERK1/2 activity and so PRC occupancy constitutes a directional molecular mechanism that synchronises neural gene accessibility and promotes engagement of the neural differentiation programme.

## Discussion

In this study, we uncover the molecular mechanism by which the neural inducing signal FGF regulates higher order chromatin organisation to orchestrate engagement of the neural differentiation programme. We demonstrate that loss of the activity of FGF-effector kinase ERK1/2 leads to rapid chromatin decompaction at the neural gene *PAX6* in both the caudal lateral epiblast of mouse embryos and in analogous hESC-derived NMP-L cells. Using ATAC-seq we show that this reflects a global action involving increased chromatin accessibility across hundreds of neural genes. Focussing on *PAX6* as an exemplar gene, we find that ERK1/2 dephosphorylation specifically results in removal of polycomb proteins, while ERK1/2 association with chromatin at this locus is unaltered. This demonstrates that ERK1/2 activity and not occupancy is critical for this mechanism. We further show that while *PAX6* is a bivalent gene that is poised to be expressed in NMP-L cells, PRC protein loss following ERK1/2 inhibition is distinct from later actions that mediate removal of the gene silencing mark H3K27me3 and transcriptional onset. Importantly, transient ERK1/2 inhibition was sufficient to trigger PRC loss and increased chromatin accessibility, indeed polycomb protein occupancy was not reinstated on resumption of ERK1/2 signalling, indicating that this is a directional molecular mechanism that advances differentiation. These findings suggest a model in which FGF/ERK1/2 signalling promotes or maintains PRCs at neural gene loci while its subsequent decline during development initiates the neural programme by synchronising removal of this repressive complex and so the increased accessibility of neural genes (Figure 7).

The ability of ERK1/2 inhibition to elicit chromatin decompaction at the *PAX6* locus in both the mouse embryo and human ESC derived NMP-L cells indicates that the molecular mechanism by which ERK1/2 regulates chromatin accessibility is conserved across species. Establishing *PAX6* as a polycomb target in both contexts further identified this repressive complex as the likely mediator of chromatin compaction. PRC1-generated compaction has been demonstrated and elucidated in mESCs [68–71] and *Pax6* has recently been identified as a PRC target in this context too [72]. The rapid decompaction of chromatin (within just one hour) following ERK1/2 inhibition in the mouse embryo shown here further suggested that ERK1/2 activity may directly maintain PRC occupancy. Exposure of differentiating human NMP-L cells to MEKi lead to loss of PRC1 component Ring1B, while PRC2 protein Jarid2 was not significantly changed. This raises the possibility that PRC1 dissociates first and is the primary target of ERK1/2 activity and PRC2 is lost secondarily. However, Jarid2 occupancy does show a marked downward trend following ERK1/2 inhibition and the lack of significant difference may reflect low amounts of Jarid2 protein at chromatin or differences in antibody effectiveness. In support of the initial loss of PRC2, we found that inhibition of ERK1/2 in NMP-L cells (which were maintained in FGF and Wnt agonist) for 3h elicited a declining trend in Jarid2 occupancy at the *PAX6* locus and that only after 12h were both Jarid2 and Ring1b significantly reduced. Importantly, in these experiments in NMP-L cells H3K27me3 levels remained unchanged and no *PAX6* transcription was detected. These findings are consistent with observations in Ring1B deficient mouse ES cells, where chromatin decompaction occurred at the *HoxB* and *D* loci that remained decorated with H3K27me3 at the gene promoters [68]. Although there are examples of transcription from loci carrying H3K27me3 [36, 73], our findings here indicate that H3K27me3 removal is closely associated with the transcriptional de-repression of *PAX6.* These findings emphasise the value of the temporal resolution afforded by this *in vitro* differentiation assay to decouple the molecular mechanisms regulating chromatin accessibility from those directing transcription initiation.

Using ATAC-seq we identified genes with increasing accessibility during neural differentiation and surveyed the global consequences of MEK inhibition in NMP-L cells in differentiation conditions; uncovering genes with increased accessibility on day 5, prior to precocious transcription of *PAX6* (on day 6). This identified hundreds of genes with GO terms associated with neural development. These included *PAX6*, but also other neural progenitor genes, (e.g. *SOX1, GlI3, GlI2, RARB, NKX6.1, DBX1, NEUROG2*, which commence expression in the embryonic body axis, like *Pax6*, after FGF/ERK1/2 signalling decline [16, 18, 31]). These data indicate that loss of ERK1/2 signalling accelerates differentiation progression by increasing chromatin accessibility at many early neural genes. The extent to which such genes are direct PRC targets or are regulated by other de-repressed neural genes remains to be determined, however, their increased accessibility on D5, prior to *PAX6* transcription, supports the possibility that they are also PRC targets.

There are a number of ways in which ERK1/2 activity may influence PRC occupancy at neural genes. Work in mES cells has shown that ERK1/2 occupies overlapping targets with polycomb proteins and in particular Jarid2 and that abrogating ERK1/2 signalling reduces Jarid2 and H3K27me3 levels raising the possibility that ERK1/2 protein is directly involved in PRC recruitment and/or maintenance [56]. Although we detected ERK1/2 at the *PAX6* locus and observed a decline in occupancy during differentiation, we also found this protein at the house-keeping gene *GAPDH*, indicating that in this context its association with DNA is not PRC target gene specific. Furthermore, inhibition of ERK1/2 activity did not alter its association with chromatin at the *PAX6* locus. This indicates that decrease in ERK1/2 activity and not ERK1/2 occupancy correlates with reduced PRC present at the *PAX6* locus in differentiating NMP-L cells. This differs from observations made in mES cells where MEKi exposure leads to an overall reduction in ERK2 occupancy at developmental genes [56]. This might reflect operation of distinct regulatory mechanisms in these different cellular contexts, indeed ERK inhibition promotes self-renewal in the mouse ES cell state, while loss of FGF and so ERK promotes differentiation progression of human ESC and of NMP-L cells. Moreover, the observation that NMP-L cells exposed to MEKi lose PRC occupancy and decompact at the *PAX6* locus, despite resumption of ERK1/2 signalling (in the continued presence of exogenous FGF), indicated that the effects of ERK1/2 inhibition are not easily reversed in NMP-L cells. This contrasts with findings in mES cells, where rescue of ERK1/2 mutant ESCs with ERK2 restored PRC mark H3K27me3 [56]. Loss of ERK1/2 signalling for a short and transient period in NMP-L cells therefore appears sufficient for sustained PRC dissociation and increased chromatin accessibility in this cell context and this provides directionality, triggering consequences which promote differentiation progression. In this sense, decline in ERK1/2 signalling here appears to be an initial step in the process of neural commitment.

In addition to potential regulation of PRC protein phosphorylation by the kinase activity of ERK1/2, a further possibility is that ERK1/2 signalling decline alters BAF (or SWI/SNF) complex composition, leading to its recruitment and so eviction of PRC proteins from gene loci, as reported in mouse fibroblasts and ESCs [74]. This is an attractive possibility as it might also provide specificity for PRC removal at neural genes, encoded in BAF complex composition [75]. This complex alters as neural progenitor cells become post-mitotic and embark on neuronal differentiation [75, 76], while here we would expect loss of ERK1/2 activity to lead to initial formation of the BAF complex characteristic of neural progenitors. In support of such a step, the transcription of key BAF complex component the ATP-dependent helicase *SMARCA2* (Brahma homolog) is sharply downregulated as FGF/ERK1/2 signalling declines in the elongating embryonic body axis [77].

The extent to which FGF/ERK1/2 signalling directs progression of differentiation by regulating PRC occupancy in contexts other than caudal epiblast/NMP-L cells requires further investigation. In mouse ESCs maintained in serum and LIF, PRC and H3K27me3 are detected at differentiation gene loci and this constitutes primed pluripotency, characterised by heterogenous cell states in a dynamic flux between pluripotency and differentiation [55, 78]. This contrasts with the ground-state or naïve pluripotent cell state achieved by treatment with both ERK1/2 and GSK3β inhibitors (2i) [25, 79], in which differentiation genes lack both PRC and H3K27me3 [55]. Importantly, differentiation of such 2i ES cells involves progression through the primed pluripotent cell state, suggesting that this involves recruitment of PRC and establishment of bivalent loci. Progression towards differentiation involves FGF/ERK1/2 signalling [28, 80] but only a transient period is required to initiate this process [27]. This last step is also consistent with the differentiation of mEpiSCs and hESCs (for which FGF is a self-renewal factor) when FGF signalling is blocked [30, 32]. The more homogenous neural differentiation achieved by expsoure of differentiaing mEpiSCs to Fgf8 [81], might also reflect more uniform recruitment of PRC. These observations together with the requirement for PRC1 and 2 for mESC differentiation [61], indicate a good correlation between exposure to a suffcient period of FGF/ERK1/2 signalling, PRC presence at differentiation genes and subsequent differentiation from the ES cell state, following ERK1/2 signalling decline.

Importantly, PRC occupancy is also associated with multiple steps that ready genes for transcription. In mouse ES cells, the presence of PRCs coincides with that of poised (phosphorylated at Ser5) RNA polymerase 2 (RNAPII), as well as with the presence of both active and repressive histone marks, as observed here for *PAX6* in NMP-L cells [34, 40, 82–85]. A role for the PRC2 associated protein Jarid2 in recruitment of poised RNAPII has also been proposed in mES cells [47]. The striking association of poised RNAPII with polycomb silenced genes in neurons, further implicates PRC occupancy with potential promoter plasticity in differentiated cells [40]. Moreover, PRC2 has been shown to mediate engagement of poised enhancers with their target genes during anterior neural induction from mESCs [86]. In addition, an in vitro interaction study has identified a role for ERK1/2-mediated phosphorylation of Ser5 on RNAPII at PRC2 occupied loci in mES cells [56]. These findings, together with our observation that ERK1/2 activity is required for PRC occupancy at *PAX6* in NMP-L cells and that its inhibition triggers precocious *PAX6* transcription in differentiation conditions, suggest that promotion of PRC at neural genes by ERK1/2 signalling functions as a rite of passage for subsequent differentiation. It will be important to investigate whether such a mechanism operates not only to maintain, but also to poise for subsequent synchronous differentiation, other FGF-expressing progenitor cell populations undergoing expansion in the developing embryo, including in limb, lung and tooth buds [87–90] as well as distinct regions of the nervous system [91]. Intriguingly, recent progress in understanding cell fate commitment during epidermal homeostasis has identified a requirement for the ERK1/2/MAPK phosphatase DUSP6 for differentiation in this adult tissue context [92], underscoring the idea that transient ERK1/2 / polycomb mediated chromatin regulation and associated gene poising constitute core and conserved molecular machinery that is a prelude to cellular differentiation.

## Materials and Methods

### Embryo dissection and hanging drop culture

CD1 mouse embryos collected at embryonic stage E8.5 were dissected in warm medium (DMEM-F12 with 10% FBS, Gibco®) and one half the litter exposed MEK inhibitor (PD184532, MEKi, 3 μM final concentration, Enzo Life Sciences) and the other half exposed to vehicle control DMSO (equal volume, 1:4000). Each embryo was placed in a drop of medium in the lid of a 35 mm plate, which was flipped over onto its base, resulting in a hanging drop [93]. Embryos were cultured in a humid chamber at 37°C and 5% CO_2_ for 1 h before fixation and processing for fluorescence in situ hybridisation (FISH) or chromatin immunoprecipitation (ChIP). All animal husbandry and procedures were approved by the UK Government Home Office and were in accordance with European Community Guidelines (directive 86/609/EEC) under project licence number 6004454.

### Cell culture of human ESCs

Human ES cells(H9, WiCell) were cultured in ES medium and provided for experiments by the Human Pluripotent Stem Cell Facility of the University of Dundee. For differentiation, cells were plated on Geltrex® (Invitrogen) coated dishes at 10^3^ cells/ cm^2^. To generate neuromesodermal progenitor-like cells (NMP-L), ES media was removed and cells were cultured with neurobasal medium (Gibco®) supplemented with N2, B27, GlutaMAX™ (final concentration for each 1x, Gibco®), 20 ng/ml bFgf (Peprotech) and 3 μM CHIR99021 (Tocris Bioscience) for 3 days (after [13]). To generate neural progenitor cells (NPs) from these NMP-Ls bFgf and CHIR99021 were removed from the medium and the cells grown for further 5 days. From day 2 to day 4 the medium was additionally supplemented with BMP and TGFβ inhibitors (Noggin, Peprotech and SB431542, Tocris Bioscience). Treatment with MEK inhibitor (PD184352, MEKi, 3 μM) or DMSO (equal volume) was administered with the culture medium in a dilution of 1:4000 for transcript level analysis and ChIP-qPCR experiments and 1:20,000 in ATAC-seq experiments.

### Fluorescence in situ hybridisation (FISH)

The mouse Pax6 fosmid pair (WIBR-1 Mouse Fosmid Library, Whitehead Institute/MIT Center for Genomic Research) and the human Pax6 BAC clone pair (WIBR-2 Human Library, see Table S3) were prepared using a standard Mini-prep protocol. Using Nick transcription the fosmids and BAC clones were labelled with Digoxigenin-11-dUTP and Biotin-16-dUTP. Unincorporated nucleotides were removed with Quick Spin G50 Sephadex columns (Roche) and labelled probes quantified by dot blotting.

Embryos were exposed for short term to MEKi or vehicle control DMSO by 1h hanging drop culture, then fixed (4% PFA, overnight), washed and dehydrated through a methanol series before being cleared in xylene and embedded in paraffin for sectioning (7 μm).

The FISH protocol on mouse tissue was adapted from (Newsome et al., 2003). Coverslips with sections containing neural tube or caudal lateral epiblast tissue were heated to 65°C for 30 min, then xylene washed (4 x 10 min each) and re-hydrated through an ethanol series to dH_2_0. The coverslips were microwaved for 20 min in 0.1 M citrate buffer, pH 6.0, then left to cool and washed in dH_2_0. For probe hybridisation, 150 ng each of the labelled probes together with 15 μg mouse Cot1 DNA and 5 μg sonicated salmon sperm DNA were denatured and incubated overnight at 37°C. After a series of washes the probes were detected using FITC conjugated anti-Digoxigenin antibody (1:20, Roche) amplified with anti-sheep Alexa Fluor 488 (1:100, Molecular Probes) and biotinylated anti-Avidin (1:100) together with Alexa streptavidin 594 (1:500, Molecular Probes); nuclei were counterstained with DAPI, coverslips were mounted with Slowfade Gold (Molecular Probes) and imaged on a Deltavision (Applied Precision).

For FISH on human ESC derived cells, cells were directly grown and differentiated on glass coverslips coated with Geltrex® and cells were untreated, exposed to DMSO or MEKi in DMSO as above. After a short fixation (10 min, 4% formaldehyde, RT) the coverslips were washed in 0.05% Triton X-100/PBS (3 x 5 min, RT). For permeabilisation the coverslips were incubated in 0.5% Triton X-100/PBS for 10 min, then transferred to 20% glycerol/PBS for 1h at 4°C and then repeatedly dipped in liquid nitrogen until completely frozen, let to thaw and soaked in 20% glycerol/PBS again (6 repeats). After the last snap freezing the coverslips were washed in 0.05% Triton X-100/PBS (3 x 5 min, RT), rinsed and incubated in 0.1 N HCl (10 min, RT) and washed in 0.05% Triton x-100/PBS (3 x 5 min, RT) again. To equilibrate, coverslips were washed in 2xSSC before pre-hybridisation (50% Formamide in 2xSSC) for 30 min at RT. For probe hybridisation, the same probe mix as for the FISH on mouse material was used (except probes specific for the human *Pax6* locus). The probes were denature on a heat block at 75°C for 3 min before hybridisation (overnight, 37°C, humid chamber). The coverslips were then washed and probes were detected using the same antibody reactions as for the FISH on mouse samples; nuclei were counterstained with DAPI, coverslips were mounted with Slowfade Gold and imaged using a widefield Deltavision Microscope.

Using OMERO insight regions of interest (ROIs) were selected over at least 3 z-sections. The 3D inter-probe distances in ROIs were measured using a custom script called OMERO mtools) [94], by segmenting the objects from the background and calculating the distance between the centroids as d in μm. For ease of comparison the ratio of number of base pairs per nm was calculated using the inter-probe distance known in bp and measured in nm.

### Chromatin immunoprecipitation (ChIP)

For each IP and control IP 25 μL of dynabeads coated with protein A (Invitrogen) were used. Beads were washed 3 times in blocking buffer (0.1% BSA in PBS) and resuspended in blocking buffer containing the antibody for immunoprecipitation or an equal amount of unspecific IgG (see Table S4) and incubated for 2 to 3h at 4°C. Beads were then washed twice in blocking buffer, resuspended in blocking buffer and stored at 4°C until immunoprecipitation.

For ChIP on human ES cells, the cells were grown and differentiated in 10 cm culture dishes at 37°C and 5% CO_2_. Cells were fixed with formaldehyde (Sigma-Aldrich®, 1%, 10 min, RT) and quenched with glycine (0.125 M, 5 min, RT). Cells were then washed 3 times with PBS and harvested by scraping into protein low bind Eppendorf® tubes and centrifuged (4000 x g, 10 min, 4°C). The pellets were snap frozen on dry ice and stored at −80°C. For the chromatin preparation, the cell pellet was resuspended in lysis buffer (50 mM Tris-HCl, pH 8.1; 10 mM EDTA, pH 8; 1% SDS and protease inhibitor), vortexted and incubated (10 min on ice). Chromatin fragments of approximately 500 to 1000 bp were generated by sonication with a probe sonicator (Vibra-Cell, Sonics®, 8 cycles, 30s on, 30s off, 30% amplitude). Sonicated lysate was diluted (1:5) in dilution buffer (20 mM Tris-HCl, pH 8.1; 150 mM NaCl; 2 mM EDTA pH 8.1% Triton X-100 and protease inhibitor) and centrifuged (20 min, full speed, 4°C). The supernatant was transferred into a fresh tube. DNA concentration was determined with NanoDrop 2000 Spectrophotometer.

For ChIP of chromatin associated proteins 100 μg of chromatin and ChIP of histone modifications 25 μg of chromatin was diluted to a final volume in 1 mL, added to the already prepared bead-antibody complexes and incubated (overnight, 4°C, rotating). An input for each ChIP (10% of ChIPed amount) was kept at −20°C. The next day, beads were washed 5 times in wash buffer (100 mM Tris, pH8.8; 0.5 M LiCl; 1% NP-40, 1% NaDoc and protease inhibitor) and then rinsed and washed in 1x TE (10 mM Tris and 1 mM EDTA, pH 8). To elute the beads were resupended in elution buffer (50 mM Tris-HCl, pH 8; 10 mM EDTA, pH 8; 1% SDS) and incubated (15 min, 65 °C, 1400 rpm). The eluate was spun down quickly and the supernatant was transferred into a fresh tube. Formaldehyde crosslinking was reversed by incubation at 65°C and 700 rpm for 6 h. The input sample was diluted with elution buffer (1:1) and crosslinking reversed. All samples were then diluted with TE (1:1) and RNA and proteins were digested (RNAse, Sigma-Aldrich®, 30 min, 37°C and proteinase K, Sigma-Aldrich®, 90 min 45°C). DNA was recovered by Phenol/Chloroform extraction (Phenol:Chloroform:Isoamylalcohol,25:24:1, pH 8, Sigma-Aldrich®) and washed with Chloroform:Isoamylalcohol (24:1, Amresco®) and then precipitated in 96% ice cold ethanol, with 3 M Na-Acetate and linear polyacrylamide (5 mg/mL, Ambion®) overnight at −80°C. Precipitated DNA was washed with 70% ice cold ethanol and pellet was air dried before resuspension in nuclease free water.

A similar protocol was used to ChIP material from caudal mouse explants. 30 caudal explants were pooled and resuspended in lysis buffer, vortexed and dissociated by pipetting before incubating on ice for 10 min and sonication. The lysate was diluted in dilution buffer, centrifuged (20 min, maximum speed, 4°C), the supernatant was transferred into a new tube (and the DNA concentration determined by NanoDrop 2000 Spectrophotometer .25 μg of chromatin were used to immunoprecipitate with 3 μg of the rabbit anti H3K27me3 antibody (Millipore) previously bound to dynabeads overnight at 4°C. After that the same procedure of washing, eluting, uncrosslinking and precipitation of DNA was used.

For quantification of specifically immunoprecipitated DNA, quantitative real time PCR was performed (PerfeCTa® SYBR® Green SuperMix for iQ™, Quanta Biosciences or Brilliant III Ultra Fast QPCR Master Mix, Agilent Technologies). Each sample was analysed in 3 biological replicates and each biological replicate was analysed in technical triplicates using primer pairs in Table S5.

### Transcription level analysis by reverse transcription quantitative PCR (RTqPCR)

For RNA extraction human ES cells were grown and differentiated in 24 well plates. Cells were washed with PBS before being collected in 350 μL of lysis buffer from the Qiagen RNeasy Mini Kit and RNA was prepared according to manufacturer’s protocol. An on-column DNA digest with RQ1 RNase-Free DNase (Promega) was performed (RT, 15 min). cDNA was generated from 500 μg purified RNA with the ImProm-II™ Reverse Transcription System (Promega) primed by random primers. The Aria Mx real time PCR system with the Brilliant III Ultra-Fast QPCR Master Mix (Agilent Technologies) was used to quantify the transcript levels. Each sample was analysed in biological triplicate and each biological replicate was run in technical triplicates (for primers see Table S6). The relative transcription level of a gene was normalised to that of *GAPDH* using the Pfaffl method [95] and *GAPDH* levels were checked by normalisation to HPRT1.

### Western Blotting

A minimum of 4 stage E8.5 mouse embryos or human ES cells grown in a 6 well plate were lysed in lysis buffer (LB Nuc+ all), kept on ice for 30 min while vortexing every 5 min and centrifugation (maximum speed, 4°C, 20 min). The supernatant was transferred into a new tube and protein concentration determined by Bradford assay. 50 μg of protein were separated on a 4-12% precast Gel (Invitrogen) in 1x MES buffer (Invitrogen) next to a size marker before they were blotted onto a PVDF membrane in transfer buffer (25 mM Tris, 190 mM glycine, 20% methanol). To minimise variances, every sample was loaded twice on the same gel and blotted onto a membrane at the same time. After blotting the membrane halved and one half was processed with a pan antibody and an antibody for the loading control α-Tubulin and the other half with a phosphorylation specific antibody and the loading control antibody. Membranes were first blocked in 3% BSA in TBST for at least 2h at RT, before incubation with primary antibody (see Table S7) overnight. After washing in TBST (3 x 10 min, RT) the membrane was incubated with secondary antibodies (see TableS7, 90 min, RT). Following TBST washes signals were visualised by scanning on LI-COR Odyssey® system. Band intensity measurements were performed for quantification. The loading controls were used to normalise for minor differences in loading before the phosphorylated ERK1/2 or PKB levels were compared to the pan ERK1/2 and PKB levels of the same samples.

### ATAC-seq

#### Experimental procedure

ATAC seq was performed on NMP-L cells, D5 and D6 cells in untreated, DMSO treated or MEKi treated condition and NP (D8) cells with the following modifications to the method described by Buenrostro and colleagues [65, 96]. Cells were treated with TrypLE™ Select (Thermo Fisher) to obtain a single cell suspension. Cells were then resuspended and counted in ice cold PBS and 50 000 cells per sample were used for the ATAC protocol (biological replicates were collected from independent experiments). Cells were pelleted and resuspended in lysis buffer (10mM Tris-HCl pH 7.4, 10mM NaCl, 3mM MgCl_2_, 0.1% NP-40) and centrifuged (10 min, 4°C). Nuclei extracts were resuspended in transposition buffer for 4 h at 37°C. The reaction was purified using the Qiagen MinElute PCR Purification kit according to manufacturer’s instructions. To generate single-indexed libraries the transposed DNA was PCR amplified with Nextera primers [65]. Library quality was assessed using Agilent TapeStation D5000 High Sensitivity Screen Tapes or a High Sensitivity DNA analysis kit for the Bioanalyzer.

#### Library-level analysis

Sequencing was performed by multiplexing 4 samples per lane on the Illumina HiSeq 2500 platform (Francis Crick Institute). This typically generated ~86 million 51bp or 101bp paired-end reads per library. Raw reads from each sample were adapter-trimmed using cutadapt (version 1.9.1) [97] with parameters “-a CTGTCTCTTATA -A CTGTCTCTTATA --minimum-length=25 –quality-cutoff=20”. If required, the additional parameters “-u <trim_length>” and “-U <trim_length>” were provided to cutadapt in order to trim all reads to 51bp. BWA (version 0.6.2) [98, 99] with default parameters was used to perform genome-wide mapping of the adapter-trimmed reads to the human hg19 genome assembly downloaded from the UCSC [100]. Read group addition, duplicate marking and insert size assessment was performed using the picard tools AddOrReplaceReadGroups, MarkDuplicates and CollectMultipleMetrics, respectively (version 2.1.1) (http://broadinstitute.github.io/picard). Reads mapped to mitochondrial DNA were removed using the pairToBed command from BEDTools (version 2.26.0-foss-2016b) [101]. Additional filtering was performed to only include uniquely mapped, properly-paired reads with insert size <= 2kb, and mismatches <= 1 in both reads. Samtools (version 1.3.1) [98] was used for bam file sorting and indexing. The filtered alignments from each library were merged at both the replicate and sample level using the picard MergeSamFiles command. Duplicate marking and removal was reperformed on the merged alignments. BedGraph coverage tracks representing the accessibility signal per million mapped reads were generated using BEDTools genomeCoverageBed with the parameters “-bg -pc –scale <SCALE_FACTOR>”. BedGraph files were converted to bigWig using the wigToBigWig binary available from the UCSC with the “-clip” parameter [102].

#### Sample-level analysis

To define peaks, regions of chromatin accessibility were identified genome-wide using MACS2 callpeak (version 2.1.1.20160309) [103] with the parameters “--gsize=hs --keep-dup all –f BAMPE --nomodel--broad”. A consensus set of intervals were obtained by merging the regions identified across all samples.

#### Replicate-level analysis

Fragment-level BED files were derived from those created using the BEDTools bamToBed command with the option “-bedpe”. Differential chromatin accessibility sites between conditions were obtained using diffReps (version 1.55.4) [104] with the parameter “—frag 0”. The annotatePeaks.pl program from HOMER (version 4.8) [105] was used to annotate the differential sites relative to the nearest gene promoter with respect to hg19 RefSeq features downloaded from the UCSC on 7th June 2017. Differential sites for each comparison were defined as those that intersected with the consensus set of accessibility sites, had an FDR <= 0.01 and fold-change >= 2, and an annotated gene symbol regardless of distance to TSS. Differential sites that were not allocated a gene symbol were not carried forward for further analysis. Genes associated with differential chromatin accessible regions as identified with ATAC-seq were used as input for gene ontology analysis using https://go.princeton.edu/.-ATAC-seq metaprofiles were generated using deepTools (version 2.5.3) [106] with the “computeMatrix scale-regions” and “plotHeatmap” commands, respectively. The genome annotation was generated by taking the minimum and maximum genomic coordinates amongst all the isoforms for a given gene. Raw ATAC-seq data and normalised replicate-level bigWig files have been deposited in the NCBI Gene Expression Omnibus (GEO) under accession code GSE121126) (access token provided to the journal editor).

## Acknowledgements

We thank Lindsay Davidson (University of Dundee, Human Pluripotent Cell Facility) for provision of quality controlled hES cells. We are grateful to Laure Verrier for sharing human NMP-L cell generation and neural differentiation protocols and her expertise in chromatin IP techniques as well as for critical discussions. We are also grateful to Peter Rugg-Gunn, Stephen Keyse, Josh Brickman, Greg Findlay and Tom Owen-Hughes as well as members of the Storey group for critical comments on versions of this manuscript. The authors thank the ENCODE Consortium and the ENCODE production labs of Bradley Bernstein, John Stamatoyannopoulos and Joseph Costello for data sets queried here. CS was supported by a BBSRC studentship (1252451) and KGS is a Wellcome Trust Investigator (WT102817AIA). JB, VM and HP are supported by the Francis Crick Institute which receives its core funding from Cancer Research UK (FC001051), the UK Medical Research Council (FC001051), and the Wellcome Trust (FC001051). We are grateful to the Francis Crick institutes’ Advanced Sequencing Facility.

## Author contributions

All experiments were carried out by CS who worked together with KGS to design experiments, analyse data and write the paper. JB VM and HP performed ATAC-seq experimental design and analysis and revised the paper.

## Supplementary Figure and Table legends

**Figure S1.**
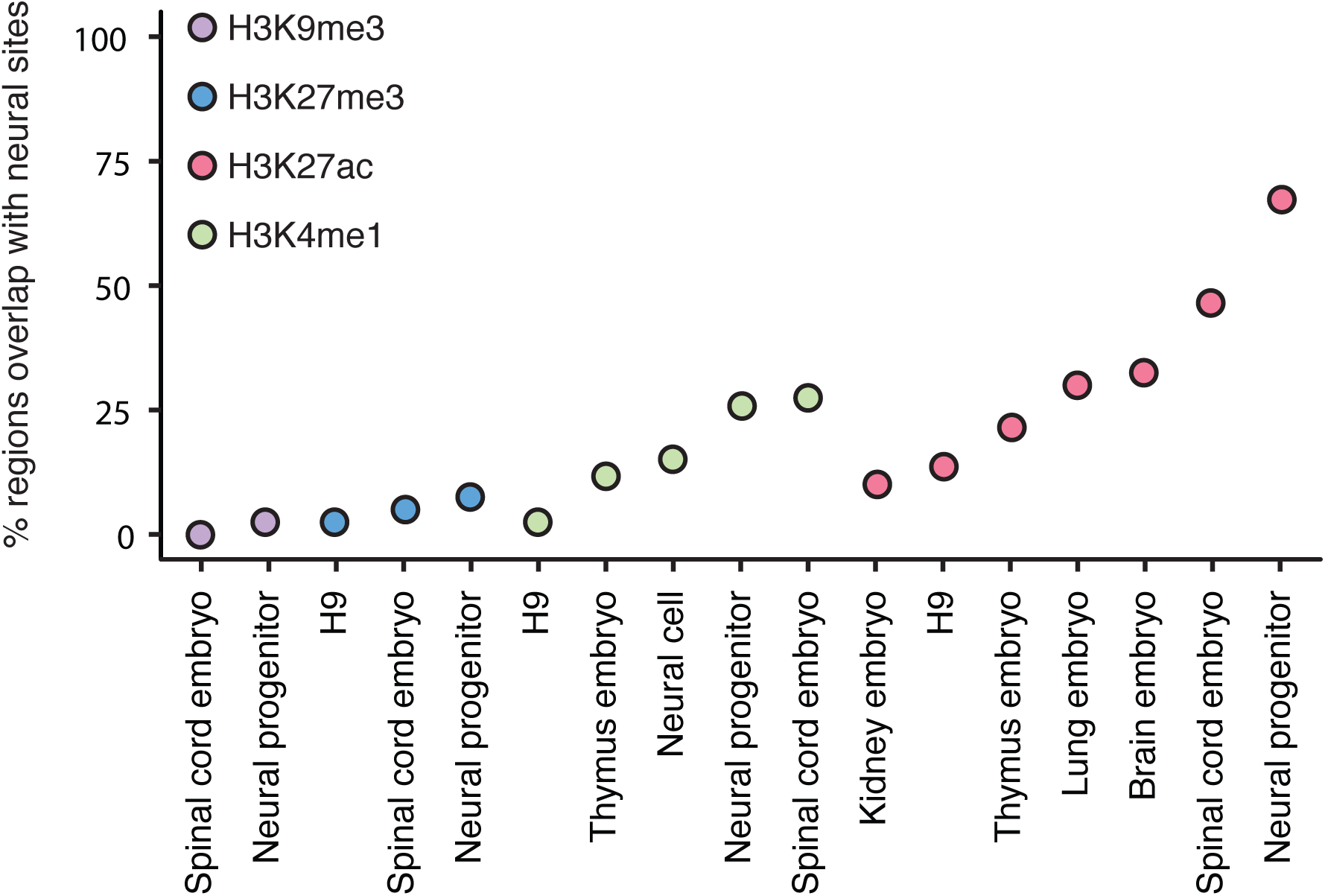
Neural sites identified by ATAC-seq overlap with known active enhancer sites in human embryonic neural tissue and *in vitro* generated neural progenitors. Comparison of peaks called from publicly available ChIPseq data sets for H3K9me3, H3K27me3, H3K4me1 and H3K27ac from the ENCODE regulatory element database (Sloan et al., 2016) with neural sites from Figure 3 show high proportion of overlap for active enhancer marks H3K4me1 and H3K27ac in neural embryonic tissue (brain and spinal cord) and *in vitro* generated neural progenitors. Data sets from embryonic lung thymus and kidney tissues were used as controls and showed smaller proportions of overlap. Furthermore H3K9me3 and H3K27me3 were used as control for repressive chromatin modification, H3K9me3 especially is known to be enriched in heterochromatin (Becker et al., 2016) and shows little to no overlap with the neural sites.

**Figure S2.**
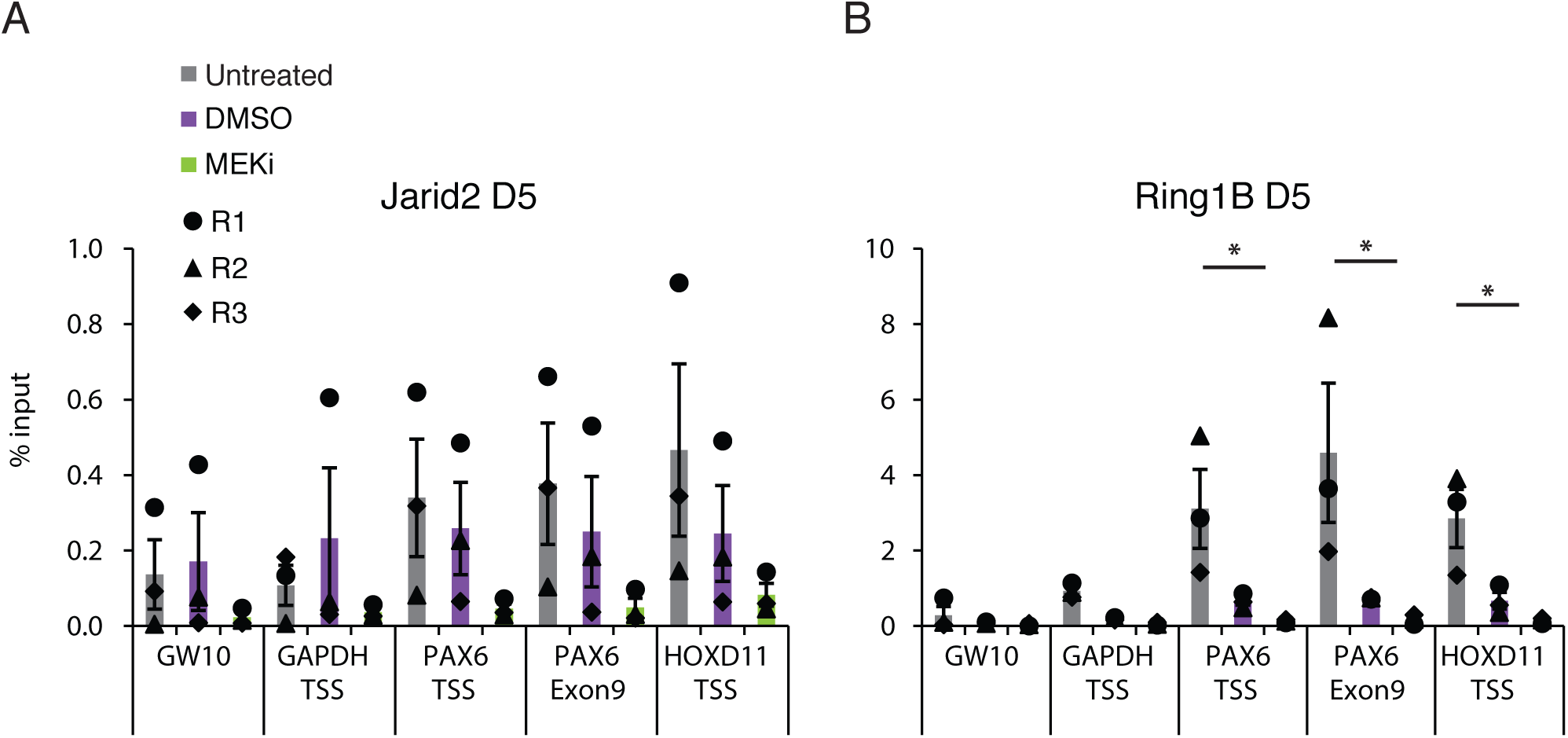
MEKi treatment does not alter PRC protein occupancy on day 5. ChIP-qPCR for Jarid2 and Ring1B for untreated, DMSO and MEKI treated cells during differentiation from NMP-L (D3) to D5 (n = 3 independent experiments indicated with circles, triangles and diamonds, bar = average, error bars = SEM, * = p ≤ 0.05 or t-test showed no significant difference);

**Figure S3.**
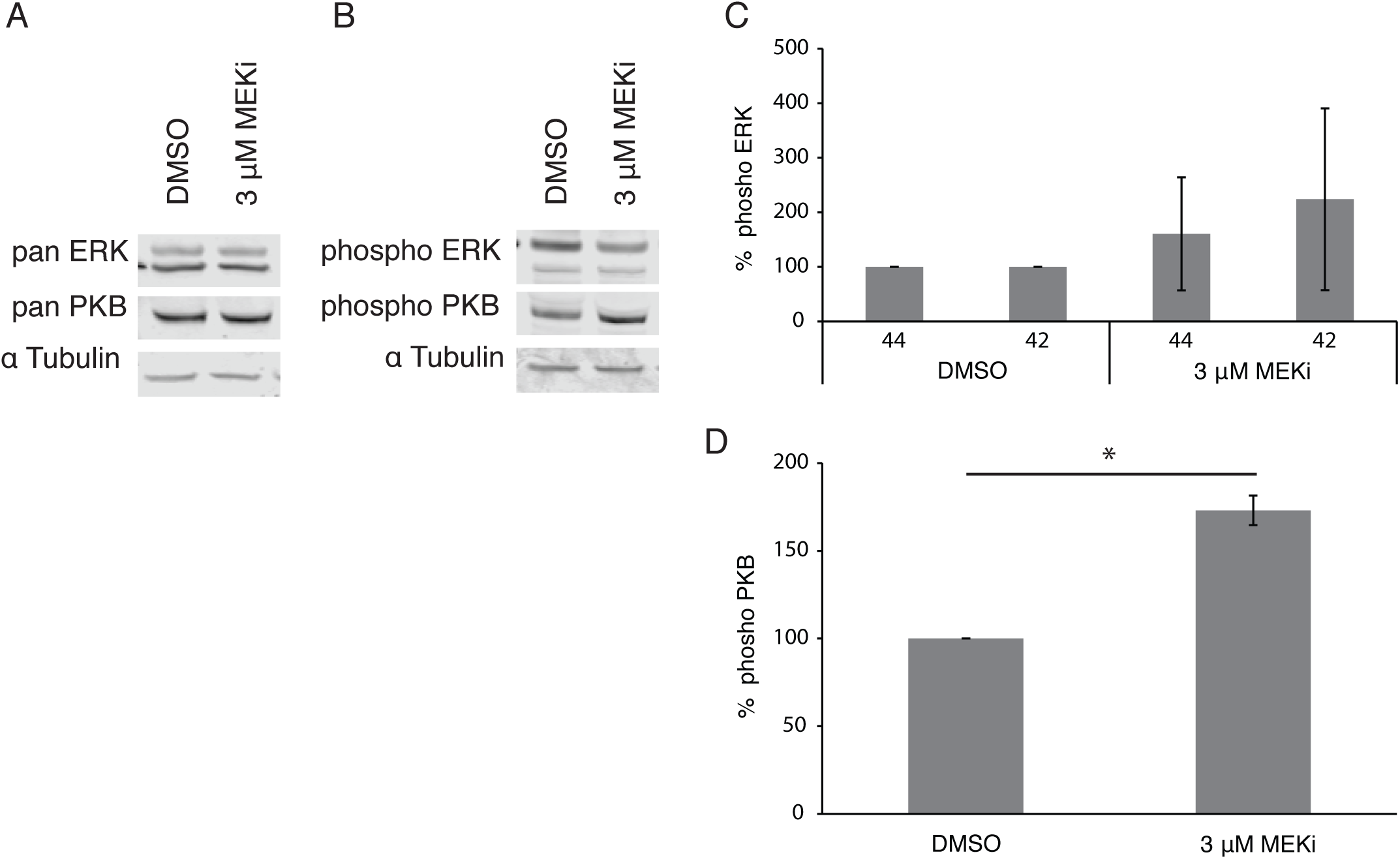
NMP-L cells cultured for 12h in MEKi no longer exhibit reduced ERK phosphorylation, but have increased PKB phosphorylation levels. (A and B) Western blot analysis of NMP-L (D3) protein extracts using phospho-specific antibodies against ERK1/2 and PKB alongside pan antibodies. (C and D) quantification of band intensity shows no reduction in ERK phosphorylation but increased levels of PKB phosphorylation (n = 3 independent experiments error bars = SEM, * = p ≤ 0.05 or t-test showed no significant difference).

**Table S1 Regions and associated genes opening during *in vitro* neural differentiation identified by ATACseq.** During *in vitro* neural differentiation from NMP-L (D3) to NP (D8) 7877 regions open up associated with 4001 genes. Opening regions are defined by chromosomal position and associated gene names have been identified by proximity.

**Table S2 List of genes associated with open chromatin regions at different time points during *in vitro* neural differentiation.** During *in vitro* neural differentiation ATAC-seq reveals that an increasing number of neural genes appear that are associated with open chromatin (by proximity). By day 5 of differentiation, regions associated with 1143 genes have opened, by day 6 regions associated with 2137 genes are open and by final day 8 regions associated with 4001 genes are now more accessible.

**Table S3 List of genes associated with open chromatin regions comparing control and MEK inhibition condition on day 5 and day 6 of the *in vitro* neural differentiation.** ATAC-seq showed that during the *in vitro* neural differentiation treatment with the MEK inhibitor PD184352 results in an increase in open chromatin. By day 5 of differentiation 1143 genes are associated with these open regions, while in the presence of MEKi, 2537 genes are associated with opened chromatin regions, 612 of which are unique to D5 MEKi condition); on day 6, 2137 genes are associated with open chromatin and with MEKi treatment regions associated with 3283 genes are open, 2045 of which were not associated with open regions in the D6 control condition and 1584 genes are unique to D6 MEKi condition.

**Table S4** Names, co-ordinates and size and interprobe distances of mouse fosmids and human BAC clones used.

**Table S5** Antibodies and amounts used for chromatin immunoprecipitation

**Table S6** Sequences of primers used for the quantification of transcript levels

**Table S7** Primary and secondary antibodies used to detect proteins transferred onto a membrane after size separation on an SDS gel.

